# Genetic variants associated mRNA stability in lung

**DOI:** 10.1101/2021.04.29.441922

**Authors:** Jian-Rong Li, Mabel Tang, Yafang Li, Christopher I Amos, Chao Cheng

## Abstract

Expression quantitative trait loci (eQTLs) analyses have been widely used to identify genetic variants associated with gene expression levels to understand what molecular mechanisms underlie genetic traits. The resultant eQTLs might affect the expression of associated genes through transcriptional or post-transcriptional regulation. In this study, we attempt to distinguish these two types of regulation by identifying genetic variants associated with mRNA stability of genes (stQTLs). Specifically, we computationally inferred mRNA stability of genes based on RNA-seq data and performed association analysis to identify stQTLs. Using the Genotype-Tissue Expression (GTEx) lung RNA-Seq data, we identified a total of 142,801 stQTLs for 3,942 genes and 186,132 eQTLs for 4,751 genes from 15,122,700 genetic variants for 13,476 genes, respectively. Interesting, our results indicated that stQTLs were enriched in the CDS and 3’UTR regions, while eQTLs are enriched in the CDS, 3’UTR, 5’UTR, and upstream regions. We also found that stQTLs are more likely than eQTLs to overlap with RNA binding protein (RBP) and microRNA (miRNA) binding sites. Our analyses demonstrate that simultaneous identification of stQTLs and eQTLs can provide more mechanistic insight on the association between genetic variants and gene expression levels.

**Author Summary:** In the past decade, many studies have identified genetic variants associated with gene expression level (eQTLs) in different phenotypes, including tissues and diseases. Gene expression is the result of cooperation between transcriptional regulation, such as transcriptional activity, and post-transcriptional regulation, such as mRNA stability. Here, we present a computational framework that take advantage of recently developed methods to estimate mRNA stability from RNA-Seq, which is widely used to estimate gene expression, and then to identify genetic variants associated with mRNA stability (stQTLs) in lung tissue. Compared to eQTLs, we found that genetic variants that affects mRNA stability are more significantly located in the CDS and 3’UTR regions, which are known to interact with RNA-binding proteins (RBPs) or microRNAs to regulate stability. In addition, stQTLs are significantly more likely to overlap the binding sites of RBPs. We show that the six RBPs that most significantly bind to stQTLs are all known to regulate mRNA stability. This pipeline of simultaneously identifying eQTLs and stQTLs using only RNA-Seq data can provide higher resolution than traditional eQTLs study to better understand the molecular mechanisms of genetic variants on the regulation of gene expression.

## Introduction

Quantitative trait loci (QTLs) are genomic loci that explain variation of a quantitative trait [1]. The most well investigated QTLs are eQTLs, which are associated with the expression level of gene transcripts [2]. Assuming different regulatory mechanisms, eQTLs proximal to and distant from the transcription start site (TSS) of genes are called *cis*-eQTLs (< 1Mb) and *trans*-eQTLs (> 5Mb), respectively [3]. By combining high-throughput gene expression data and genetic phenotype information, eQTLs can be identified systematically using a GWAS (genome-wide association study) approach [4]. It has been shown that genetic variants (single nucleotide polymorphisms) associated with complex traits, including human diseases, are more likely to be eQTLs [5]. The genetic variants located in *cis*-regulatory elements (CREs), in particular, can influence the expression of targeted genes. In fact, eQTLs are associated with many classes of CREs that are enriched in promoters, enhancers, insulators, transcription factor (TF) binding sites, and DNase hypersensitive sites (DHSs) [6–10].

Gene expression level is regulated at both the transcriptional and post-transcriptional levels. At the transcriptional level, TFs regulate the transcription rate of genes by interacting with their promoters and enhancers [11,12]. TF binding and histone modification signals in the TSS proximal regions account for over 50% of variation of gene expression [13– 15]. Genetic variants with functional impacts on TF binding motifs or promoter/enhancer accessibility are also expected to have effects on the transcription rate of related genes [16,17]. On the other hand, at the post-transcriptional level, the stability of mRNAs is under intensive regulation by microRNAs and RNA-binding proteins (RBPs) [18,19]. Genetic variants can also affect mRNA stability by interacting with microRNAs or RBPs. For example, the variant T of rs907091 located in the 3’UTR of *IZKF3* confers a miR-326 binding site, which leads to decreased mRNA stability and down-regulation of the gene; however, this is not seen with the variant C [20]. Additionally, some intronic genetic variants might also affect gene expression by interacting with splicing factors or other types of RBPs [21]. Therefore, it is often difficult to precisely interpret the eQTLs identified from high-throughput analysis. Namely, for many eQTLs, it is difficult to determine whether they influence gene expression through affecting transcriptional rate or mRNA stability. This problem is further complicated by linkage disequilibrium (LD) between neighboring genetic variants. Although high-throughput technologies that measure mRNA decay rates have been developed [22–24], there are no QTL studies that identify genetic variants associated with mRNA stability due to the lack of matched stability and genotype data.

In many eQTL studies, gene expression was determined by RNA sequencing (RNA-Seq) experiments [25–27]. Despite the protocol being designed to generate cDNA fragments from mature mRNAs, there was also a significant proportion of reads captured from intronic sequences in RNA-seq data [25]. Several studies proposed that the intronic reads of RNA-Seq were related to nascent transcription and transcriptional activity [27–30]. Based on this concept, computational methods have been developed to calculate mRNA stability based on RNA-seq data [27,31,32]. Gaidatzis et al [27] proposed a method called exon-intron split analysis (EISA) to discriminate transcriptional and post-transcriptional regulation of gene expression. Given the RNA-seq data in two experiment conditions, EISA calculates changes in reads mapped to exons (Δexon) and introns (Δintron) for each gene. It was shown that Δexon-Δintron was significantly correlated with experimentally measured mRNA stability changes between ESCs and terminal neurons. The EISA method was then further improved and then implemented in a software package, REMBRANDTS, to measure the stability of mRNAs more correctly [32].

Motivated by these methods, we developed a framework to simultaneously identify genetic variants associated with gene expression (eQTL) or mRNA stability (stQTL). We applied this framework to the lung tissue RNA-Seq data produced by the Genotype-Tissue Expression (GTEx) project [33]. For this data, we estimated the mRNA stability using REMBRANDTS and gene expression, and then performed association analysis to 15,122,700 genetic variants for 13,476 genes. We then identified a total of 186,132 eQTLs for 4,751 genes and 142,801 stQTLs for 3,942 genes. From our analysis, we found that both the stQTLs and eQTLs are enriched in the 3’UTR and CDS regions, while eQTLs are also enriched in the 5’UTR and upstream region of TSS. Compared to eQTLs, stQTLs more frequently overlapped with the binding sites of RBPs and miRNAs. To explore the role of stQTLs in mRNA stability, we took a few examples to investigate the effect of genetic variants on the binding of RBPs or TFs. Together, this study suggested that the simultaneous identification of stQTLs and eQTLs can provide a useful method to better understand the molecular mechanisms underlying genetic variants.

## Result

### Overview of this study

Fig 1 shows the rationale underlying this study. During gene expression, a gene is transcribed into a pre-mRNA, after which the introns are removed while the exons are connected into the mature mRNA. The mature mRNA is under post-transcriptional regulation by miRNAs and other mechanisms. As shown, genetic variants can not only regulate mRNA splicing but also regulate gene expression-related traits by affecting transcription rate or mRNA stability (stability QTL, denoted as stQTL hereafter). From RNA-seq data, we are able to determine the reads mapped to exonic regions to obtain gene expression levels. The mRNA stability can also be calculated by combining the reads aligned to exonic and intronic regions using the REMBRANDTS [32] algorithm. Through the eQTL analysis, genetic variants associated with gene expression are identified to obtain eQTLs. In fact, eQTLs are a mixture of QTLs that affect transcription and stQTLs, as gene expression is controlled by both transcription rate and mRNA stability. Performing an association analysis of gene expression or stability on genetic variation can identify eQTLs and stQTLs, respectively. Simultaneous identification of eQTLs and stQTLs can provide a higher resolution to understand how genetic variants affect gene expression, as well as the information to infer that a genetic variant regulates gene expression by affecting transcription activity or RNA stability. As a proof-of-concept, in this study we applied this framework to GTEx data to simultaneously investigate the eQTLs and stQTLs in lung tissue.

**Fig 1.**
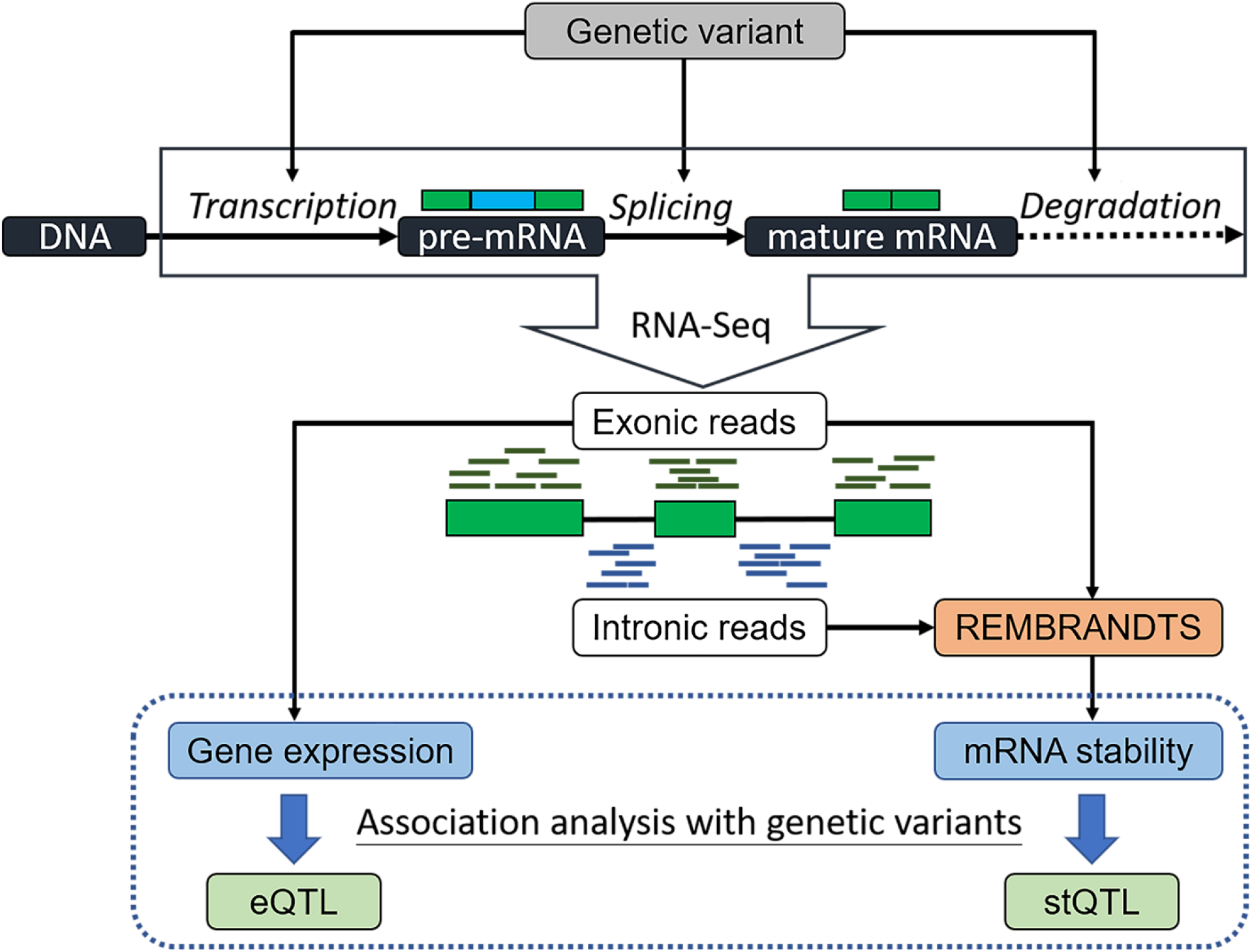
The workflow for identification of stQTL and eQTL using RNA-Seq. A genetic variant may regulate gene expression by affecting transcription, splicing, or stability at different stages of the life cycle of an mRNA. Both gene expression and mRNA stability can be estimated from RNA-Seq. Therefore, both expression quantitative trait loci (eQTLs) and stability quantitative trait loci (stQTLs) can be identified with genetic variations using the association analysis. By comparing the stQTL and eQTL, it is possible to distinguish the regulatory mechanisms underlying an eQTL.

### Expression QTLs and Stability QTLs of human lung tissue

To identify and explore stQTLs and eQTLs, we processed the raw RNA-seq data for lung tissues generated by the GTEx project. After performing quality trimming, alignment, and replicate merging from the same donors, we obtained the expression profiles of genes for a total of 289 subjects with matched genetic variation data. With REMBRANDTS, for each subject, we calculated the relative mRNA stability for 13,476 genes with intronic regions and constitutive exons. For QTL identification, we performed the association analysis on 15,122,700 genetic variants located within 100Kb upstream of TSS and 100 Kb downstream of TTS for 13,476 genes using gene expression or mRNA stability as the traits. We identified a total of 142,801 stQTLs and 186,132 eQTLs at the significance level of FDR < 5%. The numbers of QTLs were summarized in Table 1 according to the location of genetic variants on each QTL’s corresponding genes.

**Table 1.** The summary of the stQTLs and eQTLs identification in GTEx lung tissue samples. The location indicates the genomic position of the genetic variant in its corresponding gene in a QTL. The percentage was calculated from the number of QTLs at each location divided by the total number of QTLs.

| Location | Number of stQTL (%) | Number of eQTL (%) |
| --- | --- | --- |
| Upstream | 50,109 (35.09%) | 73,647 (39.57%) |
| 5'UTR | 3,877 (2.71%) | 5,802 (3.12%) |
| CDS | 3,949 (2.77%) | 4,404 (2.37%) |
| Intron | 32,675 (22.88%) | 37,232 (20.00%) |
| 3'UTR | 2,506 (1.75%) | 2,411 (1.30%) |
| Downstream | 49,685 (34.79%) | 62,636 (33.65%) |
| Total | 142,801 (100.00%) | 186,132 (100.00%) |

Ideally, we would expect that all stQTLs are also eQTLs since a genetic variant that regulates RNA stability should also affect gene expression. However, in practice, identification of different QTL types is complicated by multiple factors, including differential statistical power and LD between genetic variants. Nevertheless, we still observed that there is a very high proportion (70,105) of overlap between stQTLs and eQTLs (Fig 2A). We also found that 49% of stQTLs were also eQTLs (Fig 2B), suggesting that nearly half of stQTLs do also significantly affect gene expression. On the contrary, only 37% of eQTLs were also stQTLs. This indicated that although a considerable part of eQTLs were derived from genetic variants that significantly affect stability, more of them were regulated by genetic variants that affect other factors related to gene expression.

**Fig 2.**
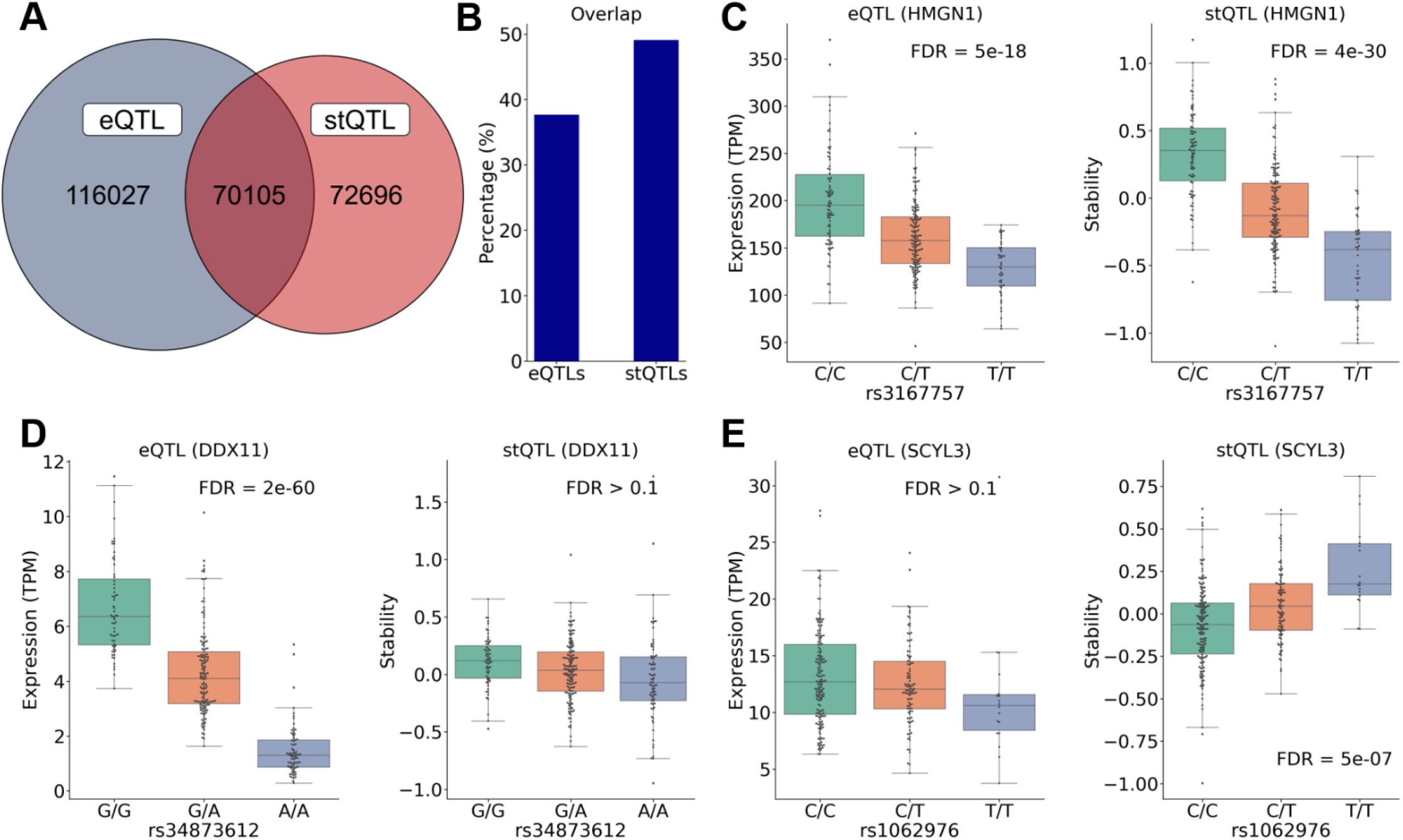
The simultaneous identification of stQTLs and eQTLs using GTEx lung tissue samples shows highly overlapped QTLs and provides additional information for investigating regulatory effects of genetic variants. (A) The Venn diagram between eQTLs and stQTLs shows that 70,105 genetic variants are both eQTLs and stQTLs. (B) The bar plot shows the percentage of the overlapped QTLs in stQTLs and eQTLs, respectively. (C) *HMGN1*-rs3167757 is an eQTL and a stQTL. The expression level and RNA stability of *HMGN1* will decrease as rs3167757 changes with CC>CT/TT. The rs3167757 is located on the binding sites of several RBPs in the 3’UTR region of *HMGN1*. (D) *DDX11*-rs34873612 is an eQTL but not a stQTL. The expression level, but not RNA stability, of the *DDX11* will decrease as rs34873612 changes with GG>GA/AA. The rs34873612 overlaps the binding sites of several TFs in 5’UTR of *DDX11*. (E) *SCYL3*-rs1062976 is an stQTL but not an eQTL. The RNA stability of the *SCYL3* will be affected by rs1062976, which is located in the 3’UTR region of *SCYL3*. The variant T of rs1062976 disrupts the binding motif of PTBP1 (destabilizer) but confers the binding motif of YBX1 (stabilizer).

By investigating stQTL and eQTL together, it is possible to determine the regulatory mechanisms underlying an eQTL. For example, genetic variant rs3167757 is significantly associated with the *HMGN1* expression level (eQTL, FDR = 5e-18) with CC>CT/TT (Fig 2C). As shown, this genetic variant is also associated with *HMGN1*’s mRNA stability (stQTL, FDR=3.7e-30). This result indicated that rs3167757 might regulate the expression level of *HMGN1* by affecting its mRNA stability. Indeed, *HMGN1*-rs3167757 has also been reported as an eQTL in lymphoblastoid cell lines (LCLs) [34,35]. The rs3167757 is located at the 3’UTR of the *HMGN1* gene and overlaps with binding sites of 20 different RBPs [36]. According to the analysis using RBPmap [37] (S1 Table), while the variant C of rs3167757 confers a motif for eight RBPs (CUG-BP, HNRNPF, MBNL1, SFPQ, TRA2B, HNRNPL, SRSF3, and YBX2), the variant T disrupts the binding motifs of five of the RBPs (HNRNPF, MBNL1, SFPQ, TRA2B, and YBX2). Notably, among them, HNRNPF [38,39], MBNL1 [40,41], and YBX2 [42] are known to contribute to mRNA stabilization. This is consistent with the observation that genotype CC is associated with higher stability of *HMGN1* mRNA than CT and TT. As another example, genetic variant rs34873612 is significantly associated with *DDX11* expression level (eQTL, FDR = 2e-60) but not with *DDX11* mRNA stability (FDR > 0.1) with GG>GA/AA (Fig 2D). This result suggested that rs34873612 might regulate the expression level of *DDX11* by affecting the transcription rate rather than its mRNA stability. According to the PROMO prediction [43], the rs34873612 is located at the 5’UTR of the *DDX11* gene and overlaps with the binding site of three TFs: GR-alpha, GATA2, and GATA3. While the variant G contributes to the binding motifs of these TFs, the variant A disrupts the binding motif of GATA3, which potentially contributes to the decreased *DDX11* expression seen in the GA and AA genotypes (Fig 2D). mRNA stability only contributes partially to gene expression level; consistently, many genetic variants are found to be stQTLs but not eQTLs. For example, rs1062976 is significantly associated with the mRNA stability of *SCYL3* (stQTL, FDR = 5e-07) but not its expression level (not an eQTL, FDR > 0.1) with CC>CT/TT (Fig 2E). Overall, our results indicated that simultaneous identification of stQTLs and eQTLs can provide us with more detailed biological insights on the regulatory effects of genetic variants on a large scale.

### Distributions of eQTLs and stQTLs across genic regions

stQTLs are associated with mRNA stability while eQTLs are associated with gene expression by affecting either mRNA stability or gene transcription. Therefore, we expect that their distributions in genes would be different. To examine this, we looked at the distribution of eQTLs and stQTLs in the DNA regions surrounding TSS and TTS of genes. We found that eQTLs are more likely to be located upstream of TSS of their corresponding genes while stQTLs tend to be located downstream of TSSs (Fig 3A). On the other hand, stQTLs are more likely to be located in the region from TTS to its 10Kb upstream than eQTLs. Both stQTLs and eQTLs are more likely to be located in the upstream region of TTS rather than the genes’ downstream regions (Fig 3B).

**Fig 3.**
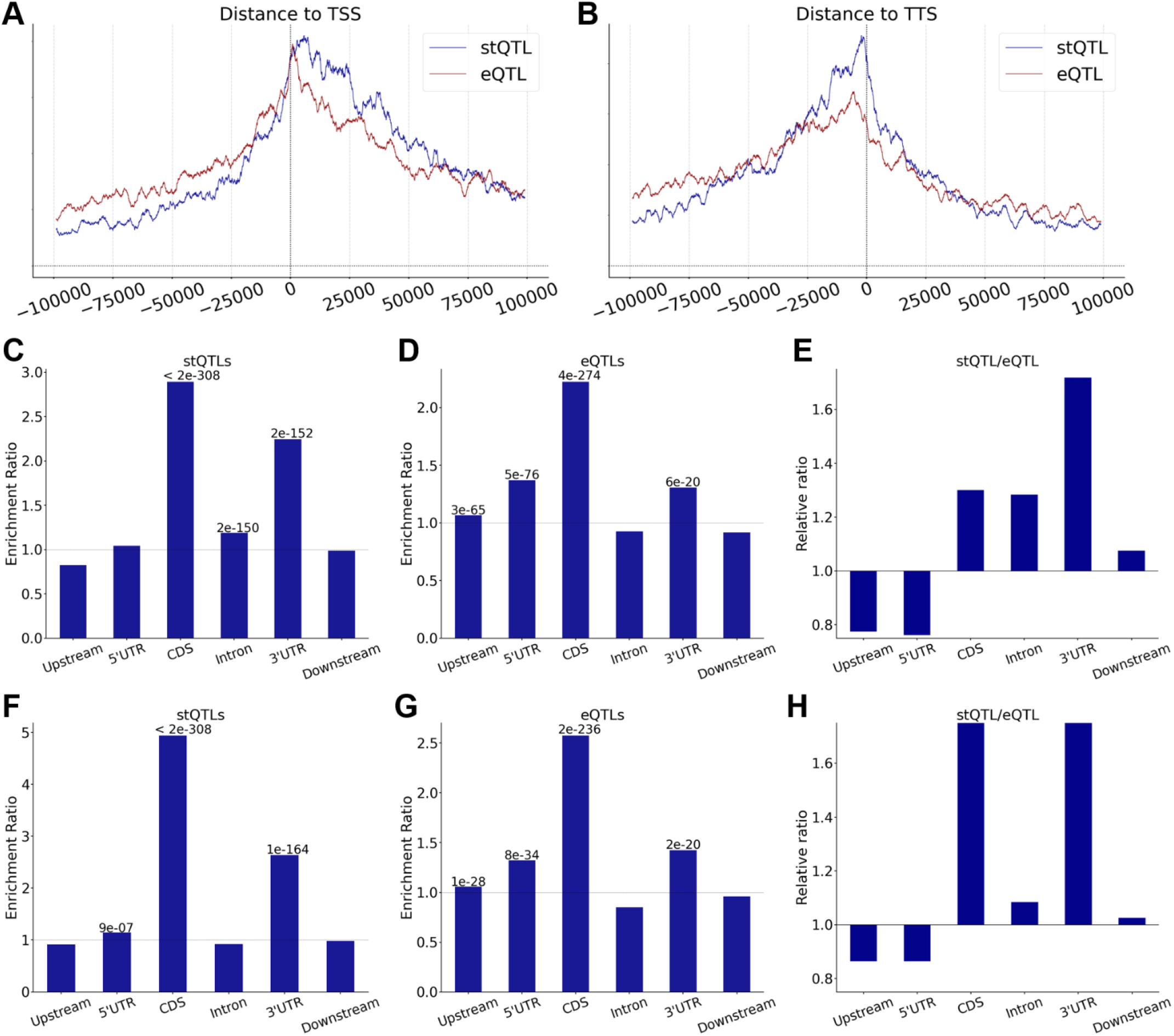
There are biased distributions in different genic regions of eQTLs and stQTLs. (A) The distribution from the enrichment ratio of stQTLs and eQTLs to TSS. Plot with the bin size of 2000 bp and the sliding window of 50 bp. (B) The distribution from the enrichment ratio of stQTLs and eQTLs to TTS. Plot with the bin size of 2000 bp and the sliding window of 50 bp. (C) The enrichment ratio in different genomic locations of stQTLs before LD filtering. The upstream indicates the region of 100 Kb upstream from TSS, and the downstream indicates the region of 100 Kb downstream from TTS. (The following figs are the same) (D) The enrichment ratio in different genomic locations of eQTLs before LD filtering. (E) The relative proportion of enrichment ratio in different genomic locations between stQTLs and eQTLs before LD filtering. (F) The enrichment ratio in different genomic locations of stQTLs after LD filtering. (G) The enrichment ratio in different genomic locations of eQTLs after LD filtering. (H) The relative proportion of enrichment ratio in different genomic locations between stQTLs and eQTLs after LD filtering.

Subsequently, we divided genomic regions associated with genes into upstream, 5’UTR, CDS, Intron, 3’UTR, and downstream regions and then examined the distributions of eQTLs and stQTLs in these regions. Using the distributions of all genetic variants as the background, we calculated the enrichment ratio of stQTLs and eQTLs by using a hypergeometric test [44]. As shown in Fig 3C, stQTLs are enriched by 2.89-fold in the CDS (P < 2e-308) and by 2.25-fold in 3’UTR (P = 2e-152). This result is consistent with the fact that genetic variants located in these regions might have functional impacts on mRNA stability by affecting RNA secondary/tertiary structure or RBP/microRNA binding. stQTLs are also slightly enriched in intron regions (ER = 1.19 and P = 2e-150). In contrast, eQTLs are enriched in the CDS (ER = 2.22, P = 4e-274), upstream (ER = 1.10, P = 3e-65), 5’UTR (ER = 1.37, P = 5e-76), and 3’UTR (ER = 1.30, P = 6e-20, Fig 3D), respectively. The enriched eQTLs in these regions may be due to the fact that gene expression can be determined not only by transcriptional activity (genetic variants in upstream, 5’UTR, or CDS regions) but also by RNA stability (genetic variants in CDS or 3’UTR regions). We compared the enrichment ratios of stQTLs and eQTLs and found that stQTLs are more likely to be located in the CDS, intron, and 3’UTR regions, while eQTLs are enriched in the upstream and 5’UTR regions (Fig 3E).

It should be noted that the resolution of QTL analysis is affected by linkage disequilibrium (LD) between neighboring genetic variants. Based on the genotype data for lung samples used in this study, we performed LD analysis and observed that many eQTL/stQTL loci were in high LD (r^2^ > 0.9) with each other (S1 Fig). To best exclude the influence of LD, we determined all LD blocks (r^2^ > 0.9) and within each block selected the most significant genetic variant as the representative stQTL/eQTL. Following that, we re-evaluated the distribution of stQTLs and eQTLs. After LD filtering, we found that the stQTLs are more enriched in CDS (ER = 4.93 and P < 2e-308), 3’UTR (ER = 2.64 and P = 1e-164), and slightly enriched at 5’UTR (ER = 1.14 and P = 9e-07). Of note, we no longer observed the enrichment of stQTLs in intron regions (Fig 3F). In contrast, the distribution of eQTLs in each gene region is similar to that before LD filtering (Fig 3G). When the distributions of stQTLs and eQTLs were directly compared, eQTLs were more likely than stQTLs to locate in the 5’UTR and upstream of genes (Fig 3H), while stQTLs are more likely to locate in the CDS and 3’UTR. No obvious difference was observed between stQTLs and eQTLs in the intron and downstream regions.

### stQTLs are significantly enriched in RBP binding sites

Having shown the enrichment of stQTLs in the 3’UTR and CDS regions, we then examined whether stQTLs tend to locate in the binding sites of RBPs or miRNAs, many of which are known to be involved in post-transcriptional regulation of mRNAs. To this end, we investigated the binding sites of RBPs and miRNAs provided by Postar2 [36] and TargetScan [45], respectively, to annotate the stQTLs identified in our analysis. Our results indicated that stQTLs (P = 3e-18, Fisher’s exact test) but not eQTLs (P > 0.1, Fisher’s exact test) are enriched in RBP binding sites. In fact, we found that 26.81% (2,770/10,332) of stQTLs overlap with the binding sites of at least one RBP, which is significantly higher (P = 7e-17, Fisher’s exact test) than 22.10% (2,788/12,617) for eQTLs (Fig 4A). In addition, we have also examined the overlap with miRNA binding sites and observed a higher proportion of stQTLs (0.19%, 20/10,332) than eQTLs (0.15%, 19/12,617) in the miRNA binding sites, although no statistical significance was detected due to very small genomic regions covered by miRNA binding sites (Fig 4B).

**Fig 4.**
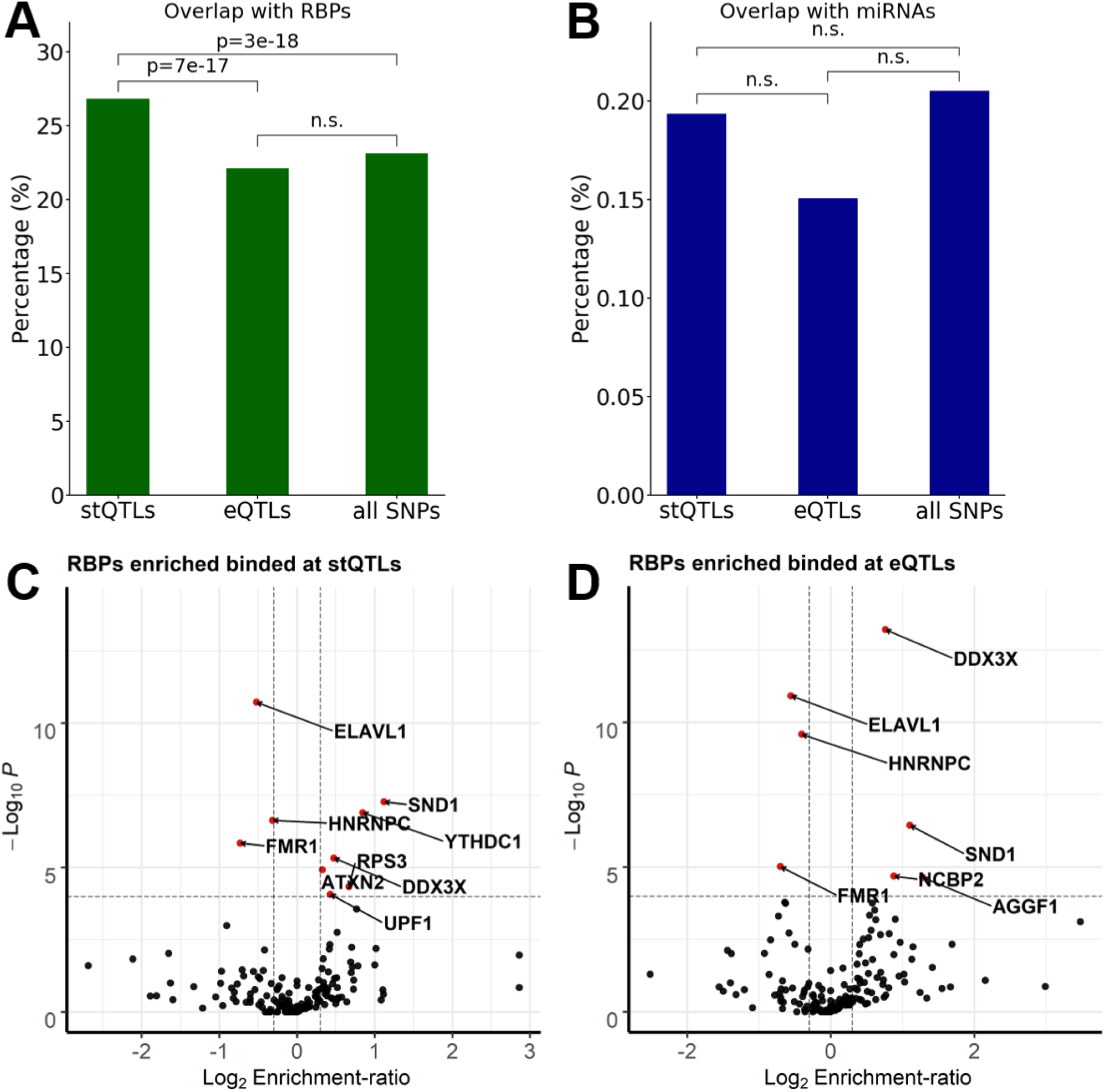
Enrichment of stQTLs and eQTLs in the binding sites of RBPs and miRNAs. (A) Proportion of overlap between stQTLs, eQTLs, and all genetic variants and RBP binding sites in mature mRNA. The statistical significance was calculated using Fisher’s exact test. The n.s. indicates not significant. (B) Proportion of overlap between stQTLs, eQTLs, and all genetic variants and miRNA binding sites in mature mRNA. The statistical significance was calculated using Fisher’s exact test. The n.s. indicates not significant. (C) The volcano plot shows that nine RBPs (red points) whose binding sites were significantly (-Log_10_ p-value > 4, 2-sides Fisher’s exact test) enriched (six RBPs, Log_2_ Enrichment-ratio > 0.3) or depleted (three RBPs, Log_2_ Enrichment-ratio < -0.3) in stQTLs. (D) The volcano plot shows that seven RBPs (red points) whose binding sites were significantly (-Log_10_ p-value > 4, 2-sides Fisher’s exact test) enriched (four RBPs, Log_2_ Enrichment-ratio > 0.3) or depleted (three RBPs, Log_2_ Enrichment-ratio < -0.3) in eQTLs.

Then we performed Fisher’s exact test to identify RBPs whose binding sites were enriched for stQTLs (S2 Table) or eQTLs (S3 Table). We identified a total of six significant RBPs (P < 1e-04) including SND1, YTHDC1, DDX3X, ATXN2, RPS3, and UPF1 (as shown in Table 2). Interestingly, SND1 [46–48], DDX3X [49,50], ATXN2 [51,52], and RPS3 [53] were known to stabilize their bound mRNAs, while UPF1 is the key factor of nonsense-mediated mRNA decay pathway [54–56]. Moreover, YTHDC1 is a well-known m^6^A (*N*^6^-Methyladenosine) reader [57], which has been found to regulate mRNA splicing [58,59], alternative polyadenylation [59], and stability [60,61] through recognizing m^6^A. Similarly, we identified four RBPs whose binding sites were significantly enriched for eQTLs (P < 1e-04, Fig 4D and Table 3), among which the two most significant RBPs, DDX3X and SND1, were also enriched for stQTLs. NCBP3 can regulate gene expression by forming a cap binding complex that binds to the 5’cap of pre-mRNA to promote splicing, 3’-end processing, and mRNA exporting [62–64], and AGGF1 was found to repress the expression of pro-inflammatory molecules [65]. These results indicate that stQTLs or eQTLs located in the binding sites of RBPs in lung tissue are indeed likely to have significant regulation on gene stability or expression by affecting the binding of the RBPs.

**Table 2.** The RBPs whose binding sites were enriched for stQTLs. Six RBPs significantly overlap (Log2 Enrichment-ratio > 0.3 and p-value < 1e-04, Fisher’s exact test) with stQTLs in mature mRNAs in lung. ER: Enrichment ratio.

| RBPs | stQTL | non-stQTL | ER | p-value | FDR |
| --- | --- | --- | --- | --- | --- |
| **SND1** | 64 | 1,504 | 2.17 | 4E-08 | 5E-06 |
| **YTHDC1** | 100 | 2,850 | 1.80 | 1E-07 | 7E-06 |
| **DDX3X** | 226 | 8,348 | 1.39 | 3E-06 | 1E-04 |
| **ATXN2** | 436 | 17,911 | 1.25 | 7E-06 | 3E-04 |
| **RPS3** | 91 | 2,923 | 1.59 | 3E-05 | 0.001 |
| **UPF1** | 198 | 7,533 | 1.35 | 5E-05 | 0.002 |

**Table 3.** The RBPs whose binding sites were enriched for eQTLs. Four RBPs significantly overlap (Log2 Enrichment ratio > 0.3 and p-value < 1e-04, Fisher’s exact test) with eQTLs in mature mRNAs in lung. ER: Enrichment ratio.

| RBPs | eQTL | non-eQTL | ER | p-value | FDR |
| --- | --- | --- | --- | --- | --- |
| **DDX3X** | 250 | 8,331 | 1.69 | 5E-14 | 1E-11 |
| **SND1** | 58 | 1,509 | 2.15 | 2E-07 | 2E-05 |
| **NCBP2** | 60 | 1,819 | 1.84 | 1E-05 | 6E-04 |
| **AGGF1** | 30 | 689 | 2.43 | 2E-05 | 6E-04 |

### Gender-specific stQTLs

We then examined whether some genetic variants were associated with mRNA stability in a gender-specific manner and denoted them as gender-specific stQTLs. We divided 289 samples into 187 males and 102 females, and then performed association analysis with covariates to implement the gender-specific stQTL classification. If a gene is specifically expressed in males or females, then an stQTL/eQTL association can only be performed in the corresponding gender. Therefore, we focused our analysis on 13,116 genes that are not differentially expressed (FDR > 0.05, t-test) between both genders and then investigated a total of 14,987,511 genetic variants located from 100Kb upstream to 100Kb downstream of a gene. Out of these gene/genetic variants, we identified 71,694 stQTLs in males and 22,841 stQTLs in females (FDR < 0.05), as well as 117,065 eQTLs in males and 48,516 eQTLs in females (FDR < 0.05), respectively. Then we defined male-specific QTLs as those that are significant in males (FDR < 0.05) but not significant in females (P > 0.1), and similarly for female-specific QTLs. In total, we identified 18,893 male-specific and 2,879 female-specific stQTLs, and 32,716 male-specific and 7,484 female-specific eQTLs. After excluding intersection with gender-specific eQTLs, we finally identified 14,683 male-specific and 2,279 female-specific stQTLs. As an example, the association between genetic variant rs397781453 and the RNA stability of *SREBP2* is female-specific (Fig 5A). As shown, we detected a significant association in females with FDR = 4e-04 but not in males (FDR ≥ 0.1). On the other hand, the association between *AQP4* and genetic variant rs12954879 is male-specific (Fig 5B). The RNA stability of *AQP4* is significantly associated (FDR = 2e-05) with genetic variant rs12954879 in males but not in females (FDR > 0.1). Of note, both *SREBP2* or *AQP4* have similar expression levels between males and females (the right panel of Fig 5A and 5B).

**Fig 5.**
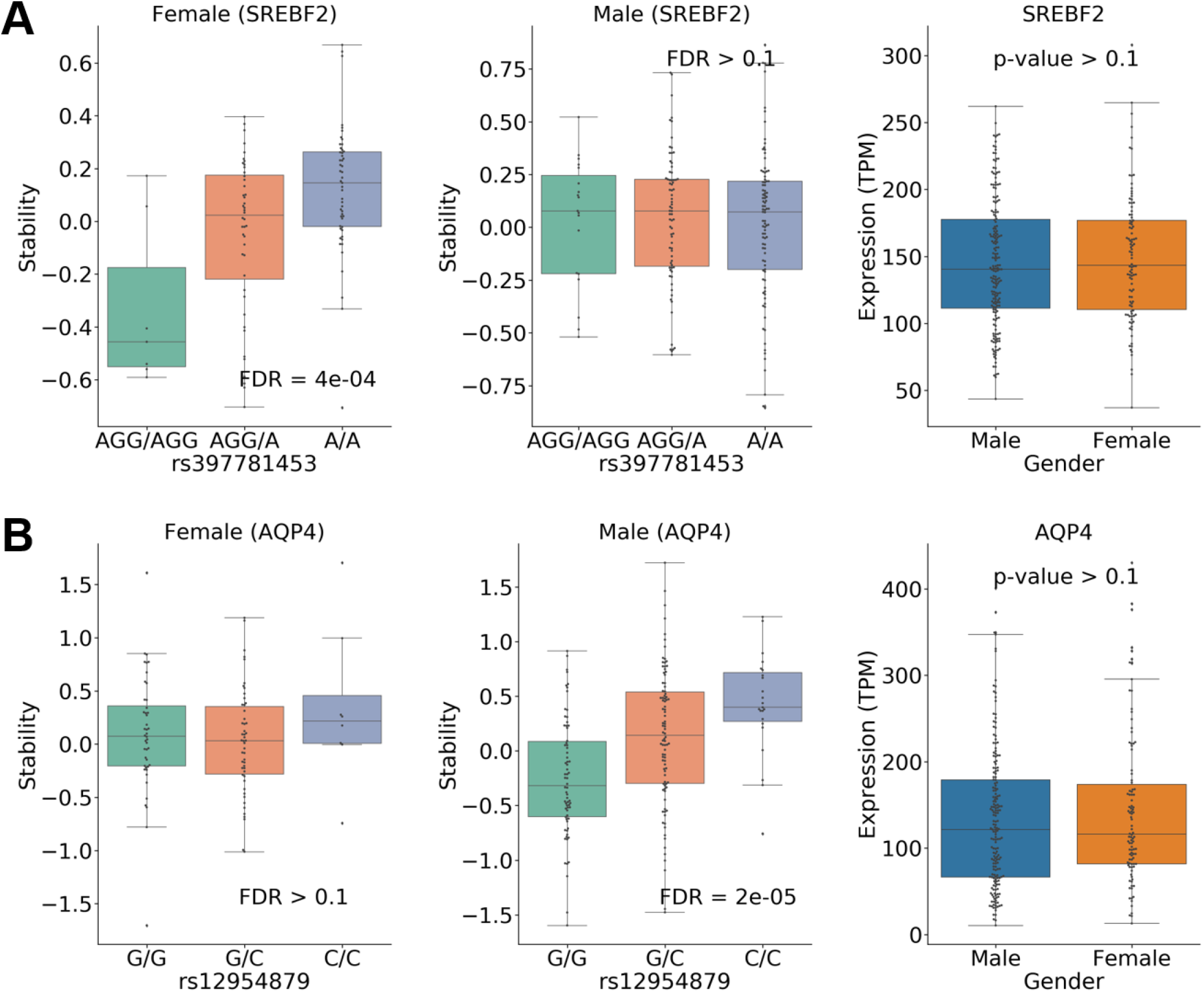
Gender-specific stQTLs identification. (A) The association between genetic variant rs397781453 and the RNA stability of *SREBP2* is female-specific (stQTL, FDR = 4e-04), but this pattern does not occur in males (FDR ≥ 0.1). In the right panel, the expression of *SREBP2* is not significantly different (P ≥ 0.1) between male and female samples. (B) The association between genetic variant rs12954879 and the RNA stability of *AQP4* is male-specific (stQTL, FDR = 2e-05), but this pattern does not occur in females (FDR ≥ 0.1). In the right panel, the expression of *AQP4* is not significantly different (P ≥ 0.1) between male and female samples.

## Discussion

In this study, we systematically identified stQTLs that are associated with mRNA stability in lung tissues and compared them with eQTLs using GTEx RNA-Seq data. Out of the 151,227,000 genetic variants within 100 Kb upstream from TSS to 100 Kb downstream from TTS of 13,476 corresponding genes, we identified a total of 186,132 eQTLs and 142,801 stQTLs. We found that stQTLs are mainly enriched in the 3’UTR and CDS regions, while eQTLs are enriched in the CDS, 5’UTR, 3’UTR, and upstream regions (Fig 3F and 3G). We also found that stQTLs are significantly located in the binding sites of RBPs (Fig 4A). Moreover, the different stQTL/eQTL variants will indeed change the motif to affect the bound RBPs, which then regulate RNA stability or gene expression (Fig 2C-2E). Our results suggest that stQTLs may significantly affect RNA stability, mostly because they are located in the 3’UTR [66,67] and CDS [68,69] regions that most often interact with other molecules. These results are consistent with previous studies, which have found that the codon usage and changes on CDS could affect its stability [68,70–72], and the sequence in 3’UTR affected mRNA stability since it includes binding sites of RBPs [73,74]. On the contrary, eQTLs are a group of complex mechanisms and may regulate expression levels by affecting stability [75,76], transcriptional activity [77–79], and even the addition of 5’cap or polyA tail [62]. Therefore, while eQTLs largely resemble stQTLs but are less enriched at the 3’ UTR and CDS regions, eQTLs are also enriched in the 5’UTR and upstream regions where the enhancers and promoters that regulate transcriptional activity are located [80–82].

Although we have shown that identifying stQTLs provides additional insights, it is worthwhile to note that determining regulatory mechanisms is largely limited by the LD between proximal genetic variants. Due to LD, it is sometimes difficult to identify the exact genetic variants that regulate gene expression. For the same reason, it is also hard to clearly distinguish genetic variants controlling gene transcription from those controlling mRNA stabilities solely based on association analysis. This analysis is expected to improve with further consideration of the location and function impact of genetic variants. In addition, the power of stQTL analysis is also limited by the computational methods used for mRNA stability inference. Although previous studies have demonstrated that the EISA algorithm [27] and its improved REMBRANDTS package [32,83,84] used in this study achieve fairly high accuracy for mRNA stability evaluation, the accuracy of inferred mRNA stability may vary significantly between different genes. First, the differential expressed long noncoding RNAs (lncRNAs) [85,86] or perturbated factors involved in intron degradation [27,87] could cause the changes of difference intronic read counts (Δintron) to affect the stability estimate. Adding the annotation of non-coding RNAs in the alignment of RNA-Seq may improve the accuracy of the mRNA stability inference. Secondly, it is difficult to accurately calculate stability for the genes with low aligned read counts because the stability inference is based on the relative change of exonic and intronic reads (Δexon–Δintron) [32]. Of note, the REMBRANDTS provides a stringency parameter to filter genes with low read counts. In our study, we set the stringency to 0.01 to include 13,429 genes for comprehensive stQTLs identification since the stability of only 2,593 genes can be calculated when the stringency is 0.9. Interestingly, we found that 41.88% (634/1,514) of stQTLs with stringency ≥ 0.9 overlap with the RBP binding sites, which is significantly higher (P = 6e-44, Fisher’s exact test) than 23.69% (1,546/6,527) of stQTLs with stringency ≤ 0.5. This result demonstrates that the stability of genes with low read counts is less accurate and can lead to potential false positives stQTLs. Finally, it should be of note that the mRNA stability calculated from RNA-Seq using REMBRANDTS is not an actual absolute value, but a differential mRNA stability relative to the average of all samples for a given gene [32,87,88]. Therefore, we suggest that it is necessary to keep these limitations in mind before evaluating mRNA stability using RNA-Seq data, and to record the stringency of the gene as a reference for the reliability of stQTLs identification.

The identification of stQTLs provides a higher resolution to better understand the molecular mechanism of genetic variants regulating gene expression, and accurate estimation of mRNA stability is very important for the identification of stQTLs. Although some high-throughput technologies, such as BRIC-Seq [24,89], have been developed to determine the decay rate of mRNA, these methods are often limited to only being used in cell culture conditions [32] and there are not enough samples available for QTLs research. Therefore, despite the limitations of computational approaches, such as SnapShot-Seq [29], EISA, and REMBRANDTS, our analysis for mRNA stability inference using RNA-seq by REMBRANDTS shows that the stQTL genic distribution and overlap with RBP binding sites is indeed consistent with biological theories. Furthermore, computer algorithms based on RNA-Seq are still under continuous development. For example, INSPEcT [87] was recently designed to calculate RNA kinetic rates based on time course RNA-seq data, or to estimate stability by calculating the difference between premature and mature RNA expression [90]. Going forward, the stQTLs which are identified with more accurate mRNA stability profiles estimation may further our understanding of how genetic variants regulate gene expression.

In conclusion, we present a large-scale identification for eQTLs and stQTLs using RNA-Seq data in lung tissues. Our results demonstrate that there are differential genic distributions as well as interactions with RBPs or TFs between eQTLs and stQTLs. We show in this study that simultaneous identification of eQTLs and stQTLs provides more biological insights for better understanding the regulatory mechanisms underlying genetic variants associated with gene expression.

## Methods

### Collection of datasets

The genotype data and RNA-Seq data of lung tissues produced by the Genotype-Tissue Expression project [33] (release 7) were used in this study. RNA-Seq SRA files and genetic variants data were downloaded from NCBI dbGaP [91] (Study Accession: phs000424.v7.p2). Subject phenotypes were collected from the GTExPortal (https://www.gtexportal.org/home/datasets). The data contains a total of 318 RNA-Seq runs and 404 genetic variant samples from 289 different subjects. We calculated the average dosages for the genetic variants data from the same subjects to represent the genotype data of subjects. For RNA-Seq analysis, the human reference genome and annotation were collected from Ensembl [92], version GRCh37.87. For RNA stability analysis, the annotation GTF files recording the coordinates of intronic and constitutive exonic segments of genes was generated using the shell script modified from the first step of the https://github.com/csglab/CRIES [32].

### Processing of RNA-Seq data

The 318 RNA-Seq SRAs were dumped into FASTQ files using SRA Toolkit (http://ncbi.github.io/sra-tools). The read quality and retained adapters were checked with FastQC [93]; then, the adapters and low-quality reads were trimmed using Trimmomatic v0.39 [94]. The alignment was performed using HISAT2 v2.1.0 [95] with default parameters. Alignment files from the same subjects were merged. Read counts of introns or exons were extracted separately using the HTSeq-count script of the HTSeq v0.12.4 [96] with the parameter --stranded=no. The RNA stability profiles for 289 subjects were estimated using the REMBRANDTS [32] with the parameter of linear method and stability stringency of 0.01. The TPM (transcripts per million) [97] was used as the expression unit to measure the expression level of 13,476 genes which have stability profiles.

### Identifying QTLs by associating genetic variants with traits derived from RNA-seq data

For covariates construction, the plink [98] (version 1.90 beta, https://www.cog-genomics.org/plink/1.9/) was performed with the parameter: --indep-pairwise 200 100 0.2 to prune a subset of genetic variants. The PCA analysis was performed after removing strand ambiguous variants (AT/CG) and genetic variants located in the MHC region. The first three PCs were selected as covariates with gender and age. For *cis*-QTL identification, genetic variants that were located within 100Kb upstream from the TSS (transcription start site) to 100Kb downstream from the TTS (transcription termination site) of Ensembl annotated genes (GRCh37.87) were selected. The expression profile was then converted with log10(TPM*100 + 1), and the linear association analysis was performed between the dosage of each genetic variant and the value of expression or stability of each gene. The Benjamini-Hochberg Procedure [99] was implemented to calculate the false discovery rate (FDR), and the genetic variant with the association of FDR less than 0.05 was regarded as a QTL.

### Enrichment analysis of QTLs in different genic regions

To determine whether eQTLs and stQTLs were evenly distributed in different genic regions, we performed the following analyses. Here we use the stQTL as an example. First, we counted the number of all genetic variants in the TSS-upstream (from TSS to 100Kb upstream), 5’UTR, CDS, 3’UTR, intronic and TSS-downstream (from TTS to 100Kb downstream) regions. Let us use *N*^*k*^ to denote the number of all genetic variants in the kth region (*k*=1, …, 6). We then counted the number of stQTLs in each of these regions and used *Q*^*k*^ to denote the number in the kth region. Third, to determine whether stQTLs are enriched in region *k*, we consider the following numbers: *Q*^*k*^, *Q*^(-*k*)^, *N*^*k*^-*Q*^*k*^, and *N*^(-*k*)^-*Q*^(-*k*)^, which (-*k*) indicate all regions other than *k*. Fisher’s exact test was then used to calculate the significance of enrichment. The enrichment was performed separately for stQTLs and eQTLs.

### Estimation of linkage disequilibrium effect

We performed the plink [98] to all genetic variants of 289 subjects with the parameter: (--r2 --ld-window 50 --ld-window-kb 100000 --ld-window-r2 0.9) to estimate the linkage disequilibrium (LD) between each genetic variant. We then constructed LD blocks, in which r^2^ of LD between each genetic variant must be greater than 0.9. To reduce the influence of LD on the gene distribution of QTLs, we selected the QTLs with the lowest FDR of the association analysis in each LD block and then performed the enrichment analysis in different genic regions as the previous section.

### Identification of QTLs located at binding sites of miRNAs or RBPs

stQTLs and eQTLs were mapped to the binding sites of RNA binding proteins (RBPs) and microRNAs (miRNAs). RBP binding site data were retrieved from Postar2 [36] (http://lulab.life.tsinghua.edu.cn/postar/). miRNA binding site data were downloaded from targetScanHuman [45] (http://www.targetscan.org/vert_72/). Both databases are based on the human genome reference version GRCh38. To match with our analysis, we performed LiftOver [100] (https://genome-store.ucsc.edu/) to convert genome coordinates into GRCh37. To evaluate QTLs that locate on the binding sites of RBPs or miRNAs, we selected stQTLs or eQTLs on mature mRNA to align the binding sites data, and then used the Fisher’s exact test [101] to identify RBPs whose binding sites were enriched located.

## Acknowledgment

All RNA-Seq data and genetic variants data used for the analyses described in this manuscript were obtained from dbGaP (accession number: phs000424.v7.p2). The phenotype data was downloaded from GTEx Portal\ (https://www.gtexportal.org/home/). This study is supported by the Cancer Prevention Research Institute of Texas (CPRIT) (RR180061 to CC) and the National Cancer Institute of the National Institutes of Health (1R21CA227996 to CC, U19CA203654 to CA). CC is a CPRIT Scholar in Cancer Research.

---

S1 Table. RBP motifs of rs3167757 variant C.

S2 Table. Fisher’s exact test between stQTL and not stQTL on mature mRNA (two-sides).

S3 Table. Fisher’s exact test between eQTL and not eQTL on mature mRNA (two-sides).

## Supporting information

**S1 Fig.**
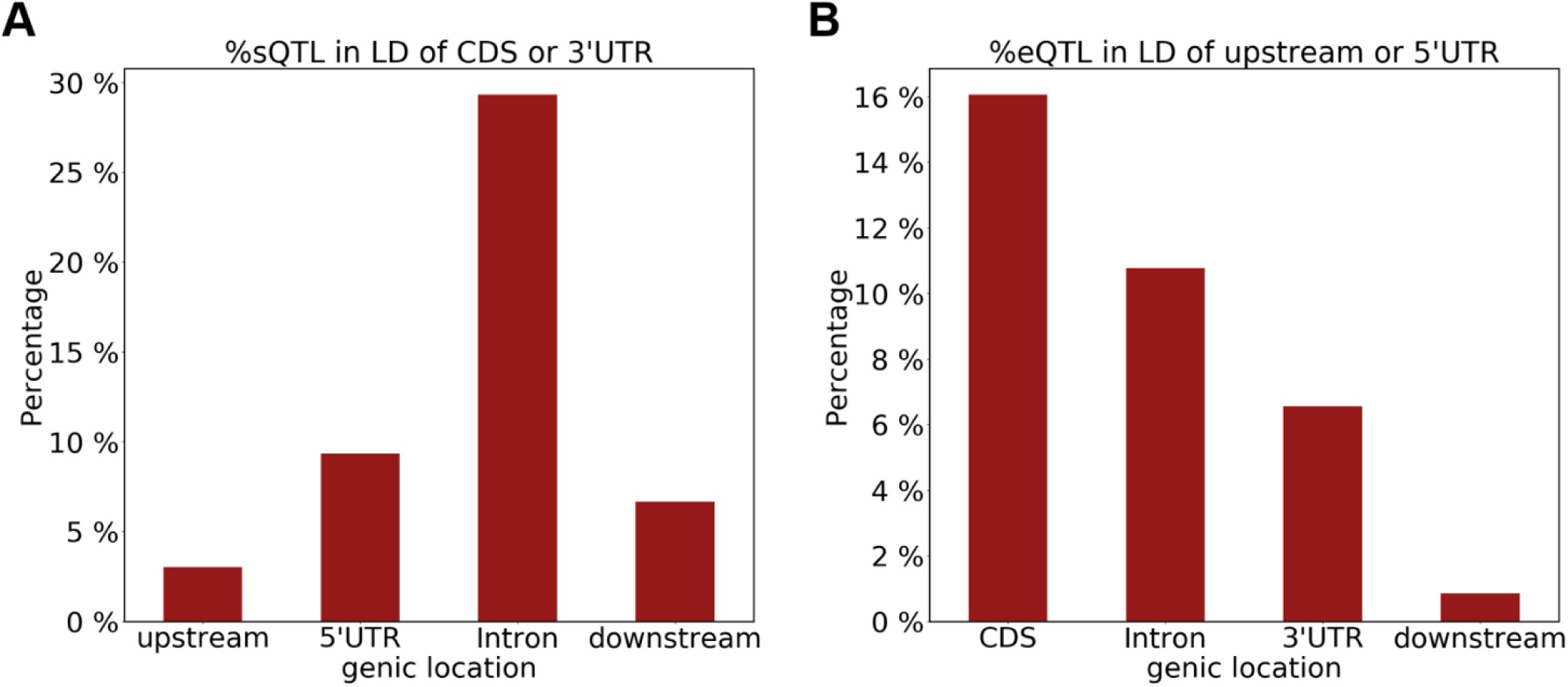
There is a considerable part of stQTLs/eQTLs have strong LD (r^**2**^> 0.9) with stQTLs/eQTLs in other regions. (A) The proportion of stQTLs in intron, 5’UTR, downstream, and upstream regions that have a strong LD with stQTLs in CDS or 3’UTR. (B) The proportion of eQTLs in CDS, intron, 3’UTR, and downstream regions that have a strong LD with eQTLs in 5’UTR or upstream regions.

